# Improved Predictions of MHC-Peptide Binding using Protein Language Models

**DOI:** 10.1101/2022.02.11.479844

**Authors:** Nasser Hashemi, Boran Hao, Mikhail Ignatov, Ioannis Paschalidis, Pirooz Vakili, Sandor Vajda, Dima Kozakov

## Abstract

Major histocompatibility complex (MHC) molecules bind to peptides from exogenous antigens, and present them on the surface of cells, allowing the immune system (T cells) to detect them. Elucidating the process of this presentation is essential for regulation and potential manipulation of the cellular immune system [1]. Predicting whether a given peptide will bind to the MHC is an important step in the above process, motivating the introduction of many computational approaches. NetMHCPan [2], a pan-specific model predicting binding of peptides to any MHC molecule, is one of the most widely used methods which focuses on solving this binary classification problem using a shallow neural network. The successful results of AI methods, especially Natural Language Processing (NLP-based) pretrained models in various applications including protein structure determination, motivated us to explore their use in this problem as well. Specifically, we considered fine-tuning these large deep learning models using as dataset the peptide-MHC sequences. Using standard metrics in this area, and the same training and test sets, we show that our model outperforms NetMHCpan4.1 which has been shown to outperform all other earlier methods [2].

## 1 Introduction

Major Histocompatibility Complex molecules (MHC) are large cell surface proteins which play a key role in immune response by detecting and responding to foreign proteins and antigens. An MHC molecule detects and binds to a peptide (a small fragment of a protein derived from an antigen), creates a peptide-MHC complex, and presents it to the surface of the cell; then, based on interactions between this complex and the T cell receptor at the cell surface, an immune response is triggered to control the compromised cell [3]. MHC molecules are classified into two classes: (i) MHC Class I which controls non-self intracellular antigens by presenting antigenic peptides (8-13 sequence length) to cytotoxic T cell lymphocytes (CD8+ TCR); (ii) MHC Class II which controls extracellular antigens by presenting antigenic peptides (13-25 sequence length) to helper T cell lymphocytes (CD4+ TCR). One of the main steps in studying the role of MHC molecules in the immune system is developing insight about the interactions of the MHC molecules and non-self pathogen peptides, referred to as MHC-peptide binding [2]. MHC-peptide binding prediction plays an important role in vaccine design and studies of infectious diseases, autoimmunity, and cancer therapy [4] [5].

There are two basic experimental methods to study MHC-peptide binding: (i) Peptide-MHC binding affinity (BA) assays in which, given a peptide, binding preferences of different MHC molecules to the peptide are calculated [6]; (ii) MHC associated eluted ligands (EL) generated by Liquid Chromatography Mass Spectrometry (LC-MS/MS) in which, based on a single experiment, a large number of eluted ligands corresponding to an MHC are identified [7]. Compared to the BA method, the EL method is highly accurate and thorough, and it is a reliable way to determine the peptides included in the immunopeptidome (namely, the whole set of peptides which have been defined in the MHC-peptides complex [8]). Both methods, on the other hand, are labor-intensive and time-consuming. As a result, a number of computational methods have been developed to predict MHC-peptide binding [9]. These started with heuristic approaches using MHC allele–specific motifs to identify potential ligands in a protein sequence [10]; later, supervised machine learning approaches were considered, including artificial neural networks (ANN) [11], hidden Markov models (HMM) [12], and regression models [13] [14]. The performance of these machine learning tools increases with the amount of data available by epitope databases such as Immune Epitope Database (IEDB) [15] and SysteMHC [16]. While some of these methods are trained for only one specific MHC allele (known as allele-specific methods), there are more generalized models (pan-specific methods) where a single model covers all of the alleles of interest in the MHC. The methods are also categorized by the type of predicted variables. Among these methods, some have been shown to be more promising, such as NetMHCPan [2], DeepLigand [17], and MHCflurry [4]. However, the most recent version of NetMHCpan (NetMHCpan 4.1) has been shown to outperform other models according to [2].

NetMHCpan is pan-specific model which predicts binding of peptides to any MHC molecule of known sequence using artificial neural networks. Since 2003, this model has gradually improved and its last version in the MHC Class I (NetMHCpan 4.1) has been introduced in 2020. This model is trained on a combination of the BA and EL peptides dataset. The inputs to this method are sequences associated with MHC-peptide complexes which are encoded by a BLOSUM matrix [18]. There are some specific features associated with this method which helps it to outperform other approaches: (i) instead of using the whole sequence of MHC molecules as input, NetMHCpan uses pseudosequences of MHC with a fixed length (34 amino acids); these pseduosequences include those amino acids associated with the binding sites of MHC which are inferred from apriori knowledge; (ii) to accommodate different lengths of peptides (8-15 in MHC Class I), they fix the length to a uniform length of 9 by insertions and deletions of amino acids associated with the peptides of different lengths; (iii) they use additional features with specificity information of the peptides during insertions and deletions steps; for example, the original length of the peptide is encoded as a categorical variable and the length of the sequence that was inserted/deleted is added as a different feature; (iv) NetMHCpan consists of several neural networks and implements the ensemble technique; in this case, using cross-validation, the training dataset is split into 5 parts and the model is trained five times, one for each split. Also, NetMHCpan uses a shallow neural network with one hidden layer which contains 56 or 66 neurons and is trained using 10 different randomly initial weight configurations; thus, the ensemble NetMHCpan contains 100 different models.

As indicated above, the most recent NetMHCpan approach (version 4.1, [2]) is based on a shallow neural network. In recent years, a number of more complex and yet efficient methods such as deep neural networks have shown promising results in a number of fields [19], [20], [21], [22], [23]. For example, transformer models, a recent breakthrough in natural language processing, have shown that large models trained on unlabeled data are able to learn powerful representations of natural languages and can lead to significant improvements in many language modeling tasks [24], [25]. On the other hand, it has been shown that collections of protein sequences can be treated as sentences so that similar techniques can be used to extract useful biological information from protein sequence databases [26], [27]. A highly successful example of this approach has been DeepMind’s recent protein-folding method, using attention-based models [28] [29] [30] [31]. Currently, there are some successful pre-trained models, publicly available, which have been shown to be helpful in a variety of downstream tasks ([32], [33], [34], [27], [26]).

Two recent works have considered using protein language models in the MHC-peptide problem. BERTMHC [35] explores whether pre-trained protein sequence models can be helpful for MHC (Class II)–peptide binding prediction by focusing on algorithms that predict the likelihood of presentation for a peptide given a set of MHC Class II molecules. They show that models generated from transfer learning, can achieve better performance on both binding and presentation prediction tasks compared to NetMHCIIpan4.0 (last version of NetMHCpan in MHC Class II [2]). Another BERT-based model known as ImmunoBERT [36] applies pre-trained transformer models in MHC Class I problem. Although they try to interpret how the BERT architecture works in MHC-peptide binding prediction, they could not compare their model fairly with NetMHCPan [2] and MHCflurry [4] due to lack of access to the same training set. Also, both BERTMHC and ImmunoBERT use the TAPE pre-trained models [26] which were trained with 31 million protein sequences, whereas now there are larger and more successful pre-trained models available such as ESM [34] and ProtTrans [32] which are trained on more than 250 million protein sequences.

In the work reported in this paper we focus on the MHC Class I peptide binding prediction and develop approaches based on the larger pre-trained protein language models; we evaluate the performance of our new model using a standard metric and the same training and test sets as NetMHCpan 4.1. We show that our methods outperform NetMHCpan 4.1 over these test sets.

## 2 Materials and Methods

### 2.1 Methods

One component of the approach in this work is based on transfer learning. In Deep Learning (DL), transfer learning is a method in which a DL model is first trained on a problem similar to the problem of interest; then, a portion or the whole of this pre-trained model is used for training the model of the desired problem. This approach is applicable when the amount of data for the problem of interest is limited, however, large databases associated with other problems with some similarity with the problem of interest exist. There are two ways to use transfer learning: (i) Fine-tuning a pre-trained model using the dataset associated with the problem of interest. In this case, a portion, or all of the weights associated with the pre-trained model are used as the initial weights of a new deep learning architecture for the desired task. (ii) Feature extraction: In this case, each input sample of the desired task is fed to the pre-trained model; then, the output or other information associated with the pre-trained model is extracted and used as features for a machine learning model. During the last decade, transfer learning has been used successfully in computer vision and more recently it has been applied to Natural Language Processing (NLP) and biology. For example, in NLP, BERT (Bidirectional Encoder Representations from Transformers) [25] is a pre-trained transformer model which is trained on a large corpus of unlabelled text including the entire Wikipedia (that’s 2,500 million words!) and the Book Corpus (800 million words). Thereafter, the pre-trained model has been used for a number of NLP tasks such as text classification, text annotation, question answering, and language inference, to name a few. BERT only uses the encoder part of the transformer since the goal is generating word embeddings (contextual relations between words (or sub-words) in a text) which are then used as features in NLP models. This method is known as self-supervision, a form of unsupervised learning in which context within the text is used to predict missing words. After BERT, various modifications based on new training methodologies and types of architecture have been attempted with the goal improving BERT (RoBERTa [37], DistilBERT [38], XLNet [39].)

Recently, following the successful results of pre-trained transformer models such as BERT and their transfer learning derivatives in NLP applications, similar approaches have been attempted in the protein field thanks to the substantial growth in the number of protein sequences. As a result, there are a number of pre-trained self-supervised BERT-like models applied to protein data in the form of unlabeled amino acid sequences which can be very useful for many protein task-specific problems using transfer learning [32] [34].

In this work we applied a fine-tuning method using two large protein language pre-trained models, Protbert-BFD [32] and ESM1b [34], two BERT-based models which are trained on hundreds of millions protein sequences. We found the performance of the ESM1b model to be slightly better than Protbert-BFD and decided to apply this model for our purposes. ESM1b is a pre-trained Transformer protein language model from Facebook AI Research [34], which has been shown to outperform all tested single-sequence protein language models across a range of protein structure prediction tasks [34]. ESM1b has 33 layers with 650 million parameters and an embedding dimension of 1280. In our work, after including an additional layer at the end of the ESM1b model, we re-trained the entire set of parameters of ESM1b and trained the parameters of the added layer using our MHC-peptide dataset. Thus, the entire architecture including the pre-trained weights of the model were updated based on our dataset (Figure 1).

**Figure 1:**
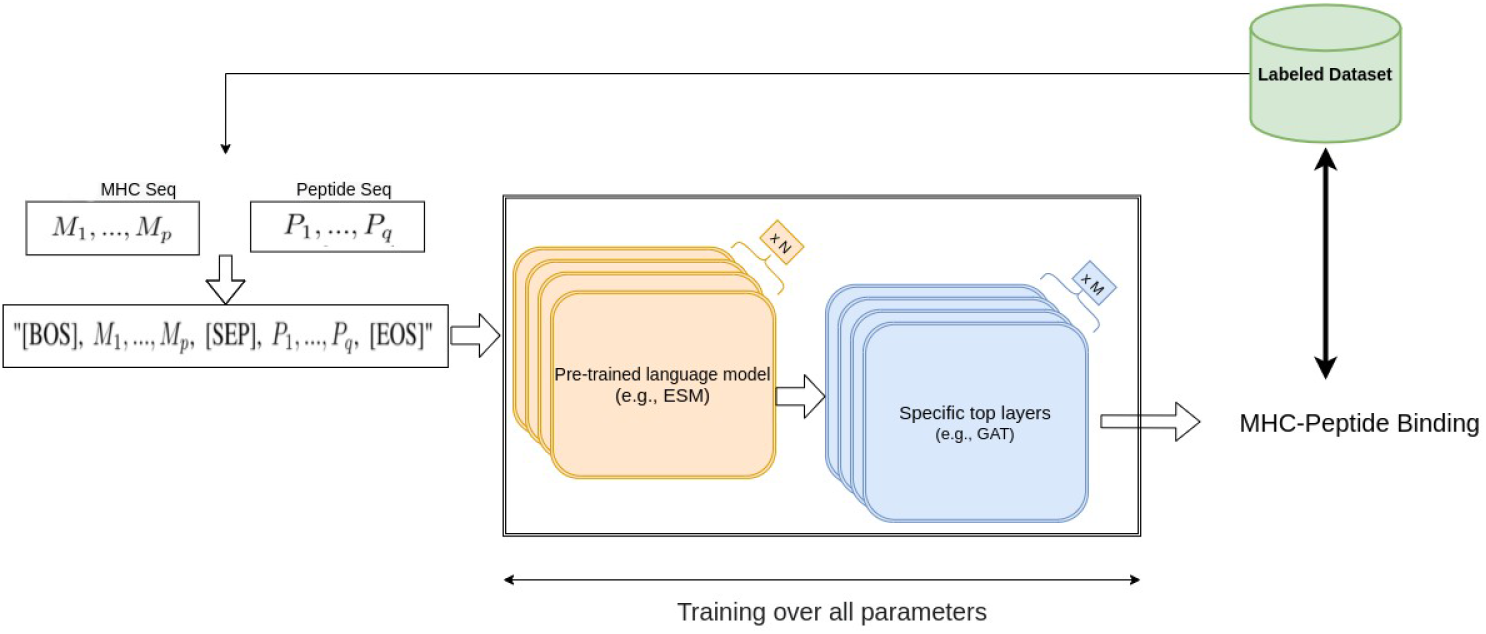
Our fine-tuning architecture based on NLP-based pre-trained models.

#### 2.1.1 ESM1b fine-tuning

Since ESM1b can be regarded as a transformer-based bidirectional language model (bi-LM), we borrowed an idea from a basic NLP task called Natural Language Inference (NLI) [40] to perform MHC-peptide binding prediction. One of the NLI tasks is the sequence-pair classification problem, namely, predicting whether a text A (e.g., “rabbits are herbivorous”) can imply the semantics in a text B (e.g., “rabbits don’t eat rats”). Similarly, in the MHC-peptide case, we would like to know whether a given peptide sequence (same as text A) binds to a given MHC sequence (same as text B), suggesting that applying an NLI-based model could be promising. A common transformer-based NLI model combines text A and B into one sequence “[BOS] seq-A [SEP] seq-B [EOS]” as input, where [BOS], [SEP] and [EOS] are special tokens ^*^ in bi-LM vocabulary.

Suppose the amino acids in the MHC and peptide sequences are *M*_1_,…, *M_p_* and *P*_1_, …,*P_q_*, respectively. We generate the sequence “[BOS], *M*_1_,…, *M_p_,* [SEP], *P*_1_,…, *P_q_*, [EOS]” with length *p* + *q* + 3 as the ESM1b input, and obtain the same size embedding vectors **v**_*BOS*_, **v**_*M*_1__,…,**v**_*M_p_*_, **v**_*SEP*_, **v**_*P*_1__,…,**v**_*P_q_*_, **v**_*EOS*_ (embedding dim 1280) from the last (33rd) layer of ESM1b, corresponding to each special token and amino acid in MHC and peptide. As a common strategy in NLP sequence classification tasks, we use the embedding of [BOS] to be the MHC-peptide sequence-pair embedding vector 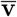. Finally, passing 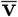 through a softmax classifier layer, we output the probability of binding and use it to compute the loss and apply back-propagation. Compared to embedding MHC and peptide separately, this compound input allows the transformer to use the attention mechanism to further extract the interactive information between the amino acids in the MHC and peptide, thus, helping the binding prediction.

Although ESM1b is well pre-trained in an unsupervised manner, using a large amount of universal sequences, we know that MHCs are a highly specific type of protein sequences, so the embedding from the pre-trained ESM1b may not be optimal for the specific MHC task and input format. Therefore, we not only need to train the final softmax classifier, but also wish to further train the ESM1b parameters to improve the sequence-pair embedding. This led us to apply fine-tuning which is commonly used in NLP. Initialized from the pre-trained ESM1b parameters, we updated the parameters in the whole network using a small learning rate during the back-propagation, so that valuable information in the pre-trained ESM1b is maintained while the fine-tuned ESM1b provided a more powerful embedding specific to the MHC tasks.

#### 2.1.2 ESM1b-GAT fine-tuning

Molecular structure-based biological data such as proteins, can be modeled with graph structure in which amino-acids or atoms are used as nodes, and contacts or bonds are used as edges. It has been shown that Graph Neural Networks (GNNs), as a branch of deep learning in non-Euclidean space, perform well in various applications in bioinformatics [41]. Here, the interaction between MHC and peptide can be described by a graph in which amino-acids are the nodes and the interaction between them can be considered as edges. To take advantage of this information, we added a variant model of GNN known as Graph Attention Network (GAT) at the top of the the ESM1b network. GAT is a novel neural network architecture that operates on graph-structured data by leveraging masked self-attentional layers to address the shortcomings of prior methods based on graph convolutions or their approximations [42]. For each MHC-peptide pair, we use a directed graph 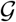, where the nodes *N*_1_,…, *N*_*p*+*q*+3_ represent the *p* + *q* + 3 tokens above, and an edge (*N_i_, N_j_*) indicates that amino acids *i* and *j* are in contact with each other. Denote the neighbor set of an amino acid *i* as 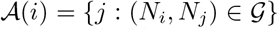; then, each embedding vector **v**_*i*_ is updated as a weighted average of its transformed neighbor embedding vectors:

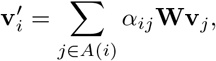

where **W** is a weight matrix for vector transformation, and the weight *α_ij_* is computed using an attention mechanism. Suppose **z**_*ij*_ is the concatenation of vectors **Wv**_*i*_, and **Wv**_*j*_ and **c** is a parameter vector, then the weight *α_ij_* is given by:

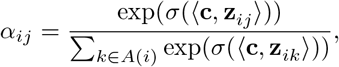

where *σ* is an activation function.

After each GAT layer, we update the embedding vector for the amino acids and the special tokens as 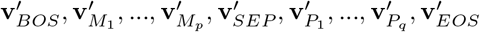, and more GAT layers follow. Here, in our implementation, we use two fully connected GAT layers. Same as vanilla transformer model [24], we apply multi-head attention mechanism in which for each GAT layer, we split the parameters and pass each split independently through a separate head. Particularly, in the first GAT layer we use 8 attention heads which are then concatenated together and passed to the next layer while in the final GAT layer we average the heads of a certain token. We finally use the embedding vector of [BOS] in the final GAT layer as the MHC-peptide sequence pair embedding vector to determine binding prediction. The final GAT layer was meant to use the attention mechanism to aggregate all the node information into [BOS] position by letting [BOS] token contact with all the amino acids in graphs, which makes the [BOS] embedding potentially a more powerful sequence embedding than simply using the average of the embedding vectors output by the first GAT layer. Compared to using only ESM1b layers, now we can introduce more prior information of contact from graphs, which will be used by the GAT layers to dynamically refine the ESM1b embedding.

### 2.2 Dataset

#### 2.2.1 Training set

We used the training set used by the last version of NetMHCpan [2], including 13 millions binary labeled MHC-peptide binding samples, generated from two main data sources (i): the BA peptides derived from in-vitro Peptide-MHC binding assays, and (ii) the EL peptides derived from mass spectrometry experiments. However, it has been shown that the results from the mass spectrometry EL experiment are mostly poly-specific, i.e., they contain ligands matching multiple binding motifs [8]. That being said, for most of the samples in the EL dataset, each peptide is associated with multiple alleles (from 2 to 6 alleles for each peptide). Thus, in this training set, the EL dataset is composed of two subsets: (i): Single-Allele (SA, peptides assigned to single MHCs) and (ii) Multi-Allele (MA, peptides with multiple MHC options to be assigned). Table 1 shows the distribution of the aforementioned dataset which indicates that more than 67% of the dataset is associated with EL-MA. According to [8], the existence of the MA dataset introduces some challenges in terms of data analysis and interpretation; therefore, to train a binary MHC-peptide predictor, a process, known as deconvoluting the MA binding motifs, is needed to convert these EL-MA data to a single peptide-MHC pair [2].

**Table 1:** Distribution of training set used in NetMHCpan 4.1 [2]; Columns correspond to each type of training data, for which the number of positive and negative samples, and the total amount of unique MHCs are shown. A threshold of 500 nM is used to define positive BA data points.

| Binding Affinity |  |  | EL (Single Allele) |  |  | EL (Multi Allele) |  |  |
| --- | --- | --- | --- | --- | --- | --- | --- | --- |
| positives | Negatives | MHCs | positives | Negatives | MHCs | positives | Negatives | MHCs |
| 52,402 | 155,691 | 170 | 218,962 | 3,813,877 | 142 | 446,530 | 8,395,021 | 112 |

#### 2.2.2 Deconvolution of Multi Allelic (MA) data

To deconvolute the EL-MA dataset, several computational approaches have been used based on unsupervised sequence clustering [43] [44]. Although these methods show some progress in dealing with the MA dataset, they have some shortcomings; for example, they do not work in cell lines including MHC alleles with similar binding motifs. Therefore, in the new version of NetMHCPan (Version 4.1), they present a new framework, NNAlign-MA [8], which works better than the previous approaches. NNAlign-MA is a neural network framework, which is able to deconvolute the MA dataset during the training of the MHC-peptide binding predictor. Recently, [35] attempted to solve this problem in MHC Class II by using a multiple instance learning (MIL) framework. MIL is a supervised machine learning approach, where the task is to learn from data including positive and negative bags of instances. Each bag may contain many instances and a bag is labeled positive if at least one instance in it is positive [45]. Assume the ith bag includes m alleles as *A_i_* = {*a*_*i*1_, *a*_*i*2_,…, *a_im_*} which is associated with peptide sequence *s_i_*. At each training epoch, for each instance in the ith bag, *x_ij_* = (*a_ij_, s_i_*), the probability of whether that instance is positive, *p*(*y_ij_* = 1|*x_ij_*) is defined as 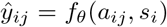 where *f_θ_* is the neural network model; then, in [35], they use max pooling as a symmetric pooling operator to calculate the prediction of the bag from the predictions of instances within it. Here, in our work, we follow this MIL idea to deal with the EL-MA dataset.

#### 2.2.3 Test set

In order to have a fair comparison of our model and NetMHCPan 4.1, we used the same test set they provided in their work (Table 2). This dataset is associated with a collection of 36 EL-SA datasets, downloaded from [46]. Each dataset is well enriched, length-wise, with a number of negative decoy peptides equal to 5 times the number of ligands of the most abundant peptide length.

**Table 2:**
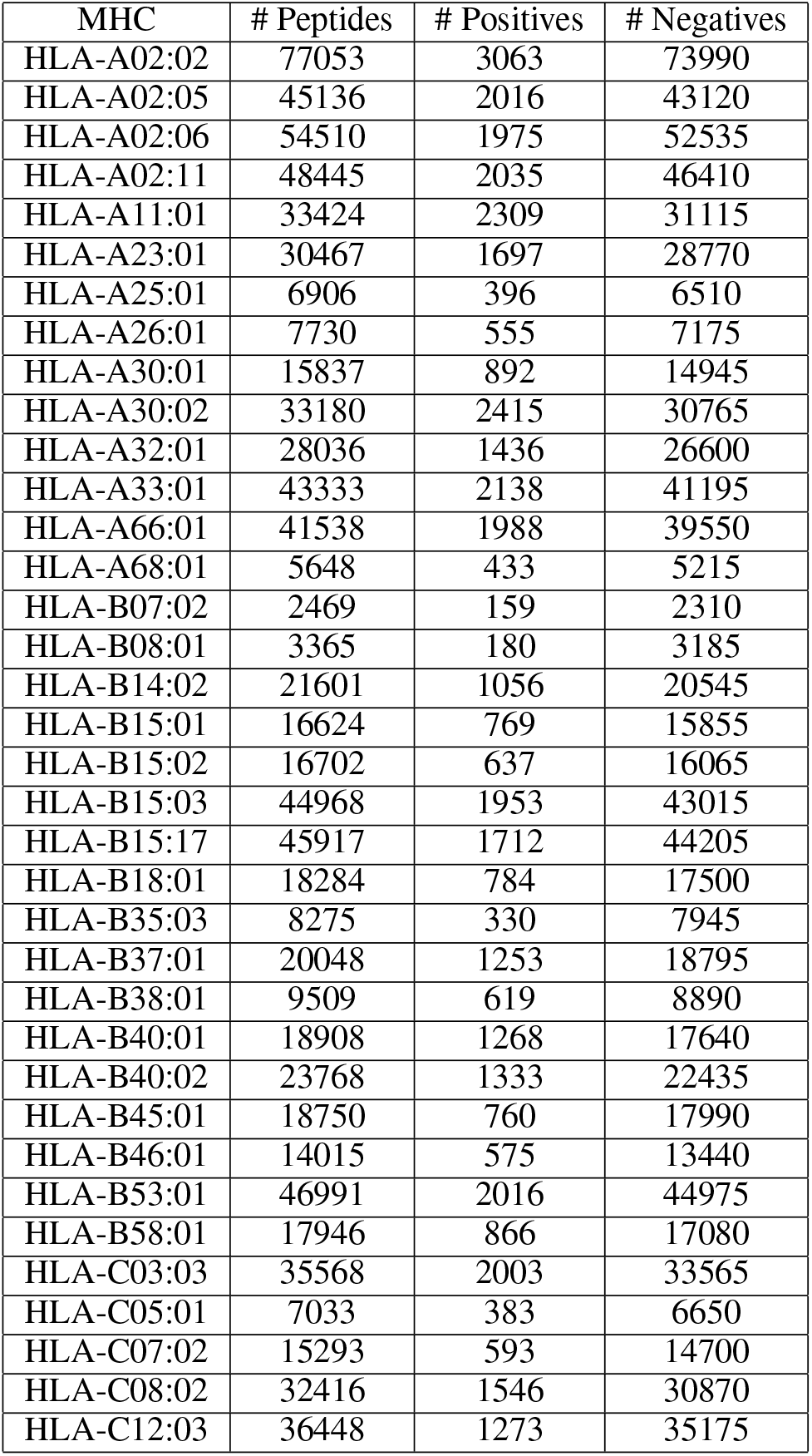
Independent EL SA test set provided by NetMHCpan 4.1 ([2]

| MHC | # Peptides | # Positives | # Negatives |
| --- | --- | --- | --- |
| HLA-A02:02 | 77053 | 3063 | 73990 |
| HLA-A02:05 | 45136 | 2016 | 43120 |
| HLA-A02:06 | 54510 | 1975 | 52535 |
| HLA-A02:11 | 48445 | 2035 | 46410 |
| HLA-A11:01 | 33424 | 2309 | 31115 |
| HLA-A23:01 | 30467 | 1697 | 28770 |
| HLA-A25:01 | 6906 | 396 | 6510 |
| HLA-A26:01 | 7730 | 555 | 7175 |
| HLA-A30:01 | 15837 | 892 | 14945 |
| HLA-A30:02 | 33180 | 2415 | 30765 |
| HLA-A32:01 | 28036 | 1436 | 26600 |
| HLA-A33:01 | 43333 | 2138 | 41195 |
| HLA-A66:01 | 41538 | 1988 | 39550 |
| HLA-A68:01 | 5648 | 433 | 5215 |
| HLA-B07:02 | 2469 | 159 | 2310 |
| HLA-B08:01 | 3365 | 180 | 3185 |
| HLA-B14:02 | 21601 | 1056 | 20545 |
| HLA-B15:01 | 16624 | 769 | 15855 |
| HLA-B15:02 | 16702 | 637 | 16065 |
| HLA-B15:03 | 44968 | 1953 | 43015 |
| HLA-B15:17 | 45917 | 1712 | 44205 |
| HLA-B18:01 | 18284 | 784 | 17500 |
| HLA-B35:03 | 8275 | 330 | 7945 |
| HLA-B37:01 | 20048 | 1253 | 18795 |
| HLA-B38:01 | 9509 | 619 | 8890 |
| HLA-B40:01 | 18908 | 1268 | 17640 |
| HLA-B40:02 | 23768 | 1333 | 22435 |
| HLA-B45:01 | 18750 | 760 | 17990 |
| HLA-B46:01 | 14015 | 575 | 13440 |
| HLA-B53:01 | 46991 | 2016 | 44975 |
| HLA-B58:01 | 17946 | 866 | 17080 |
| HLA-C03:03 | 35568 | 2003 | 33565 |
| HLA-C05:01 | 7033 | 383 | 6650 |
| HLA-C07:02 | 15293 | 593 | 14700 |
| HLA-C08:02 | 32416 | 1546 | 30870 |
| HLA-C12:03 | 36448 | 1273 | 35175 |

### 2.3 Metric

Predicting the binding affinity of MHC with a peptide is a binary classification problem. Typical metrics for assessing the quality of binary classification models for a given task include precision, accuracy, recall, receiver operating characteristic curve (ROC) and the corresponding Area Under the Curve (AUC). In this work, we use AUC and a specific precision metrics known as positive predictive value (PPV); AUC and PPV have been used as the main metrics in previous work in MHC-peptide binding prediction [2] [4]. AUC is an evaluation metric for binary classification problems which measures the area underneath the receiver operating characteristic curve (ROC). AUC ranges in value from 0 to 1 and models with higher AUC perform better at distinguishing between the positive and negative classes. To calculate AUC, we use Scikit-learn, a free software machine learning library for Python programming language. PPV is another metric which specifically is defined in this area and is interpretable as a model’s ability to rank positive samples far above the negative samples. PPV is defined as fraction of true positive samples (hits) among the top-scoring 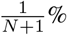 samples, provided that ratio of the number of negative samples (decoys) to positive is N:1 (N is known as hit-decoy ratio). Since NetMHCpan [2] uses hit-ratio 19 and MHCflurry [4] uses N=99, here in this work, we use 19, 49 and 99.

## 3 Results

Using the above test set, we calculated the AUC and PPV scores of our ESM fine-tuning method. In order to evaluate and compare our performance with the state-of-the-art methods, we used the latest version of NetMHCpan server (Version 4.1) which, according to their studies ([2]), and using the same training and test sets, outperformed other methods including MHCflurry [47], MHCFlurry-EL, and MixMHCpred [44]. We used three different hit-decoy ratios (19, 49 and 99) for PPV calculations. Table 3 shows that using the AUC metric, our method works better than NetMHCpan. In addition, as seen in Figures 2, 3 and 4, our model outperforms NetMHCpan over all hit-decoy ratios in the 35 different test sets; only for HL-B18:01, NetMHCpan performs better.

**Table 3:** Comparison of AUC between our model and NetMHCpan (V4.1) ([2]

| Allele | Our model AUC | NetMHCpan4.1 AUC |
| --- | --- | --- |
| HLA-A02:02 | 0.99 | 0.98 |
| HLA-A02:05 | 0.98 | 0.95 |
| HLA-A02:06 | 0.99 | 0.98 |
| HLA-A02:11 | 0.98 | 0.97 |
| HLA-A11:01 | 0.96 | 0.95 |
| HLA-A23:01 | 0.97 | 0.90 |
| HLA-A25:01 | 0.98 | 0.94 |
| HLA-A26:01 | 0.97 | 0.93 |
| HLA-A30:01 | 0.98 | 0.95 |
| HLA-A30:02 | 0.97 | 0.96 |
| HLA-A32:01 | 0.98 | 0.97 |
| HLA-A33:01 | 0.99 | 0.98 |
| HLA-A66:01 | 0.99 | 0.98 |
| HLA-A68:01 | 0.96 | 0.91 |
| HLA-B07:02 | 0.97 | 0.89 |
| HLA-B08:01 | 0.98 | 0.95 |
| HLA-B14:02 | 0.98 | 0.96 |
| HLA-B15:01 | 0.97 | 0.94 |
| HLA-B15:02 | 0.97 | 0.95 |
| HLA-B15:03 | 0.99 | 0.98 |
| HLA-B15:17 | 0.99 | 0.98 |
| HLA-B18:01 | 0.98 | 0.96 |
| HLA-B35:03 | 0.98 | 0.95 |
| HLA-B37:01 | 0.97 | 0.92 |
| HLA-B38:01 | 0.98 | 0.94 |
| HLA-B40:01 | 0.99 | 0.98 |
| HLA-B40:02 | 0.98 | 0.97 |
| HLA-B45:01 | 0.99 | 0.97 |
| HLA-B46:01 | 0.98 | 0.95 |
| HLA-B53:01 | 0.99 | 0.99 |
| HLA-B58:01 | 0.98 | 0.95 |
| HLA-C03:03 | 0.90 | 0.79 |
| HLA-C05:01 | 0.98 | 0.92 |
| HLA-C07:02 | 0.98 | 0.97 |
| HLA-C08:02 | 0.99 | 0.96 |
| HLA-C12:03 | 0.98 | 0.97 |

**Figure 2:**
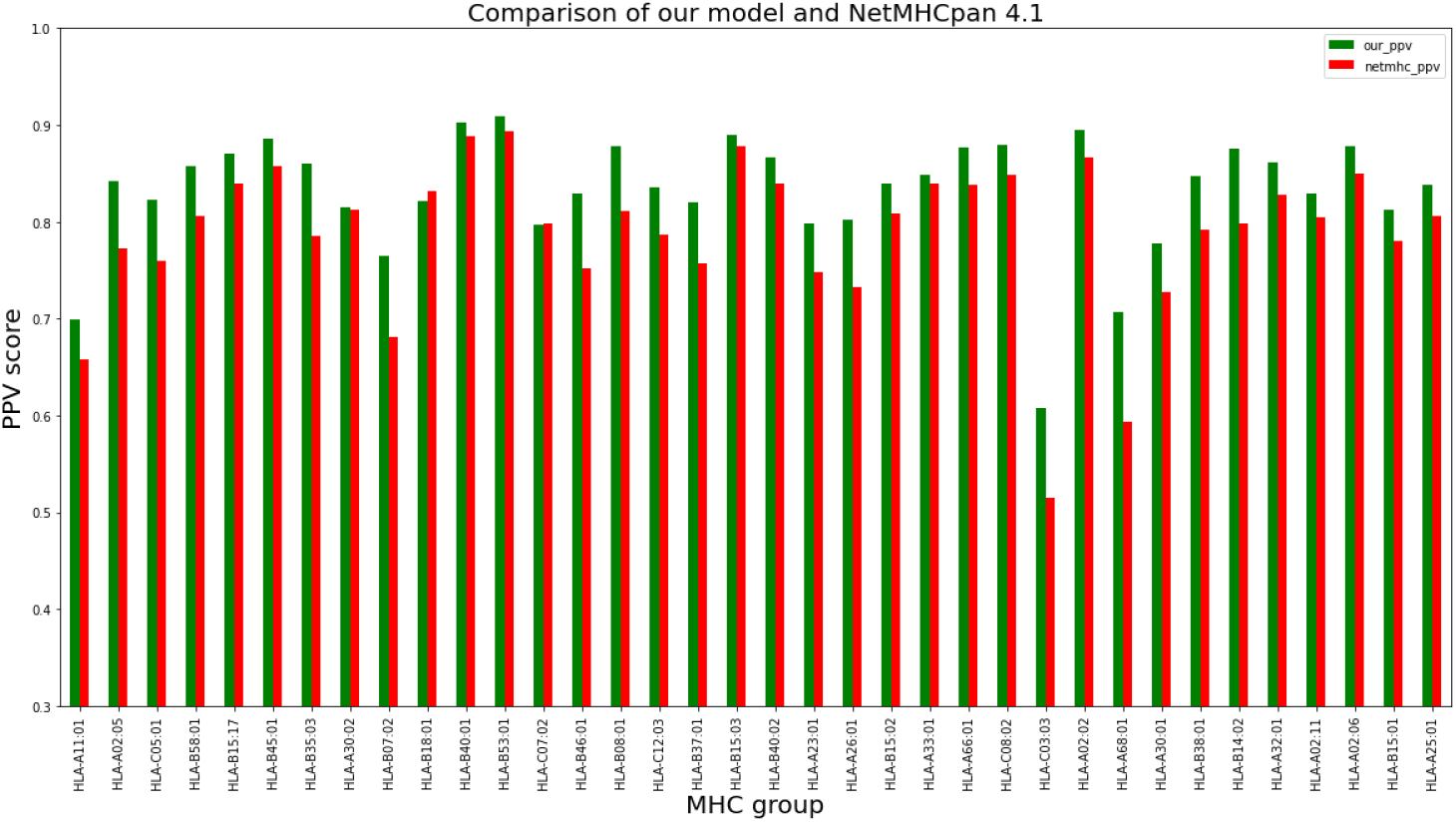
PPV Comparison (hit-decoy ratio: 1:19) of our ESM fine-tuning method with the latest NetMHCpan server (Version 4.1) over the same training and test sets [2].

**Figure 3:**
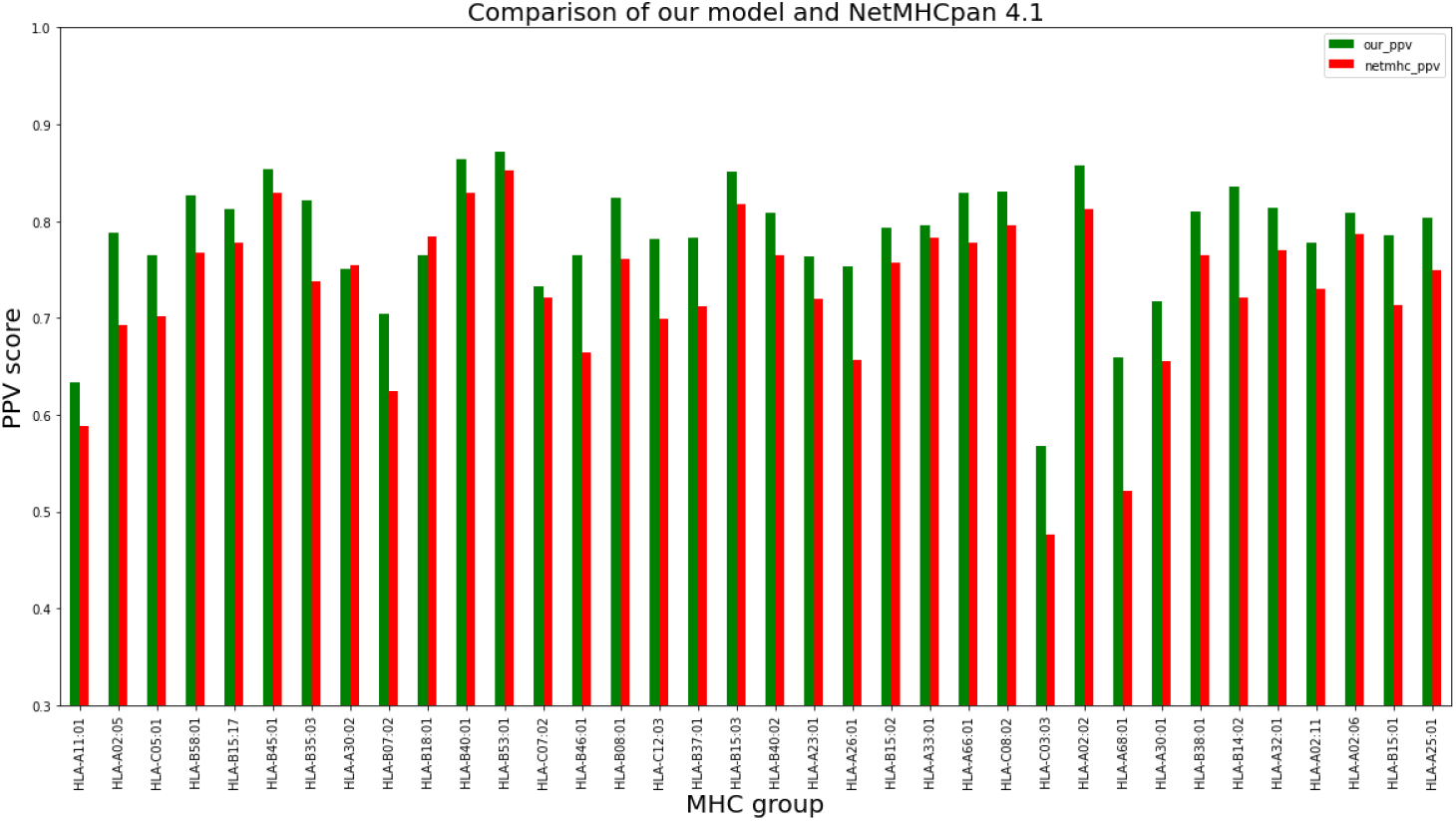
PPV Comparison (hit-decoy ratio: 1:49) of our ESM fine-tuning method with the latest NetMHCpan server (Version 4.1) over the same training and test sets [2].

**Figure 4:**
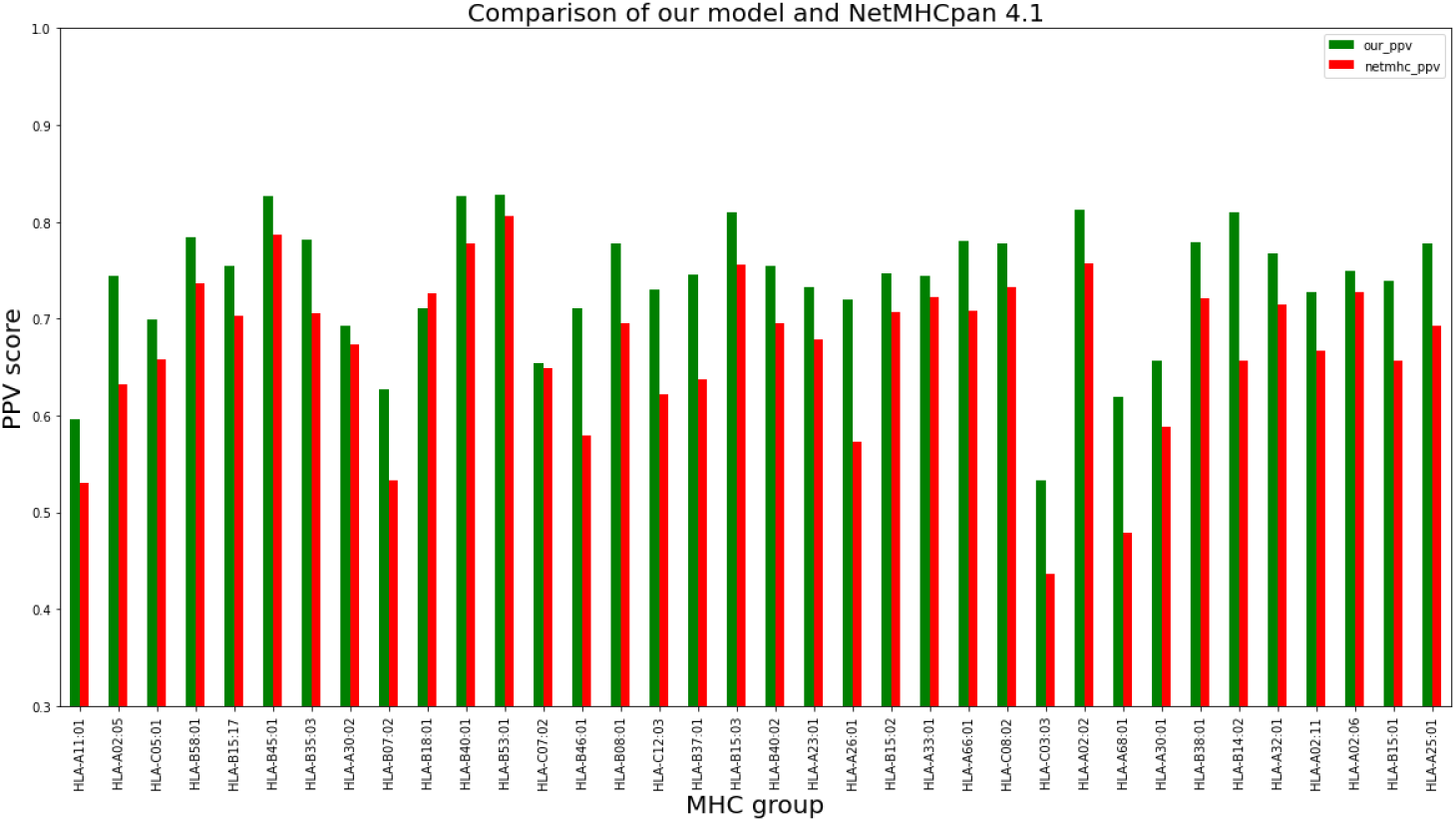
PPV Comparison (hit-decoy ratio: 1:99) of our ESM fine-tuning method with the latest NetMHCpan server (Version 4.1) over the same training and test sets [2].

To compare the GAT-ESM fine-tuning method versus the vanilla ESM, we use the subsets of training sets that include samples associated with peptides of length 8 and 9 and compare both methods over the test set. As can be seen in Figure 5, ESM-GAT significantly outperforms the ESM method when the test set with peptide length 10-15 is considered, but the results are almost the same when using the test set with peptides of length 8 and 9 (Figure 6). It follows that GAT improves the ability of the model to predict the peptides with lengths different from those considered in the training set, which might be useful for training models beyond MHC Type I.

**Figure 5:**
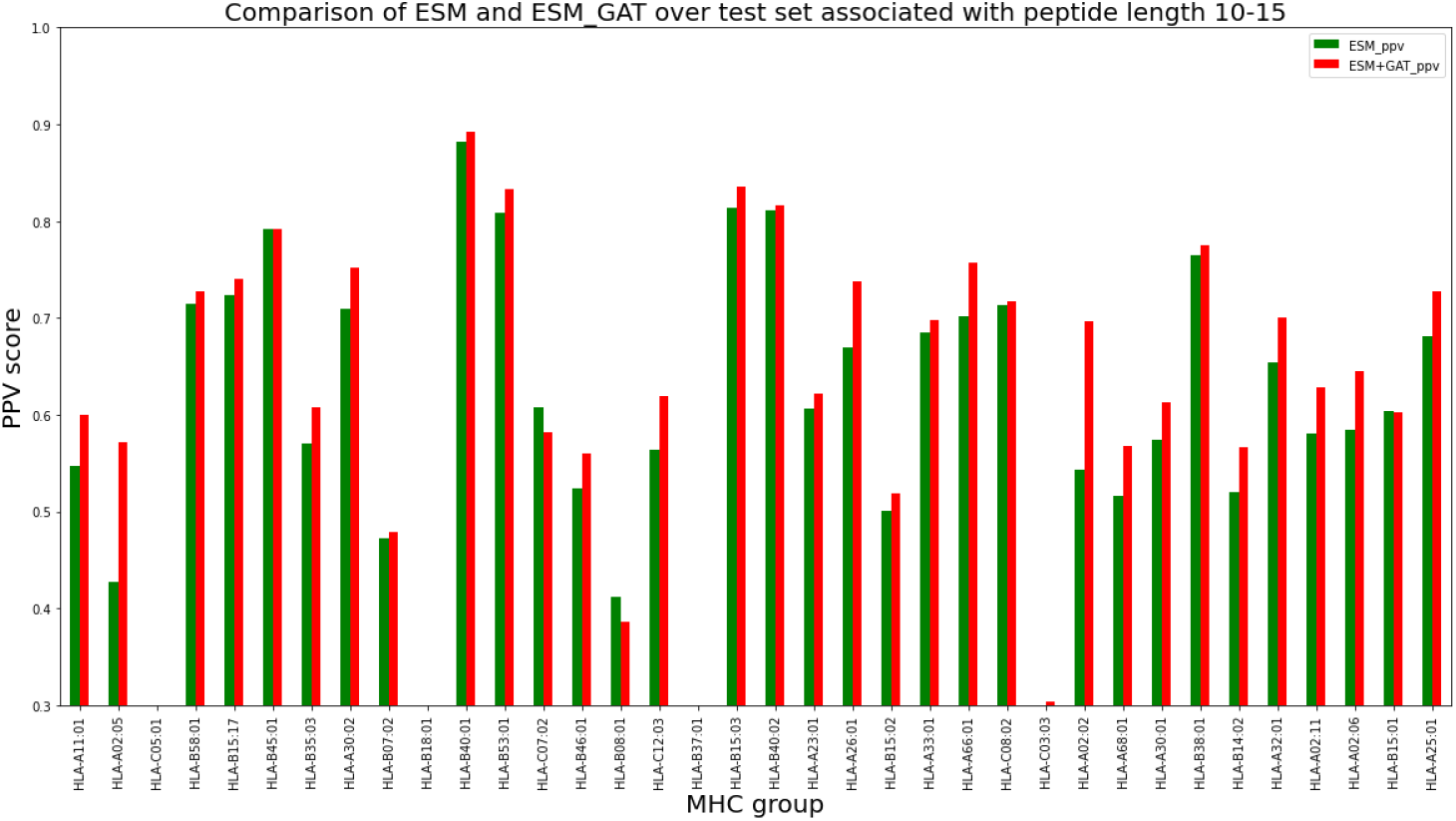
PPV Comparison (hit-decoy ratio: 1:19) of ESM fine-tuning method versus ESM-GAT over the training set with peptide length 8 and 9 and test set with peptide length 10 to 15.

**Figure 6:**
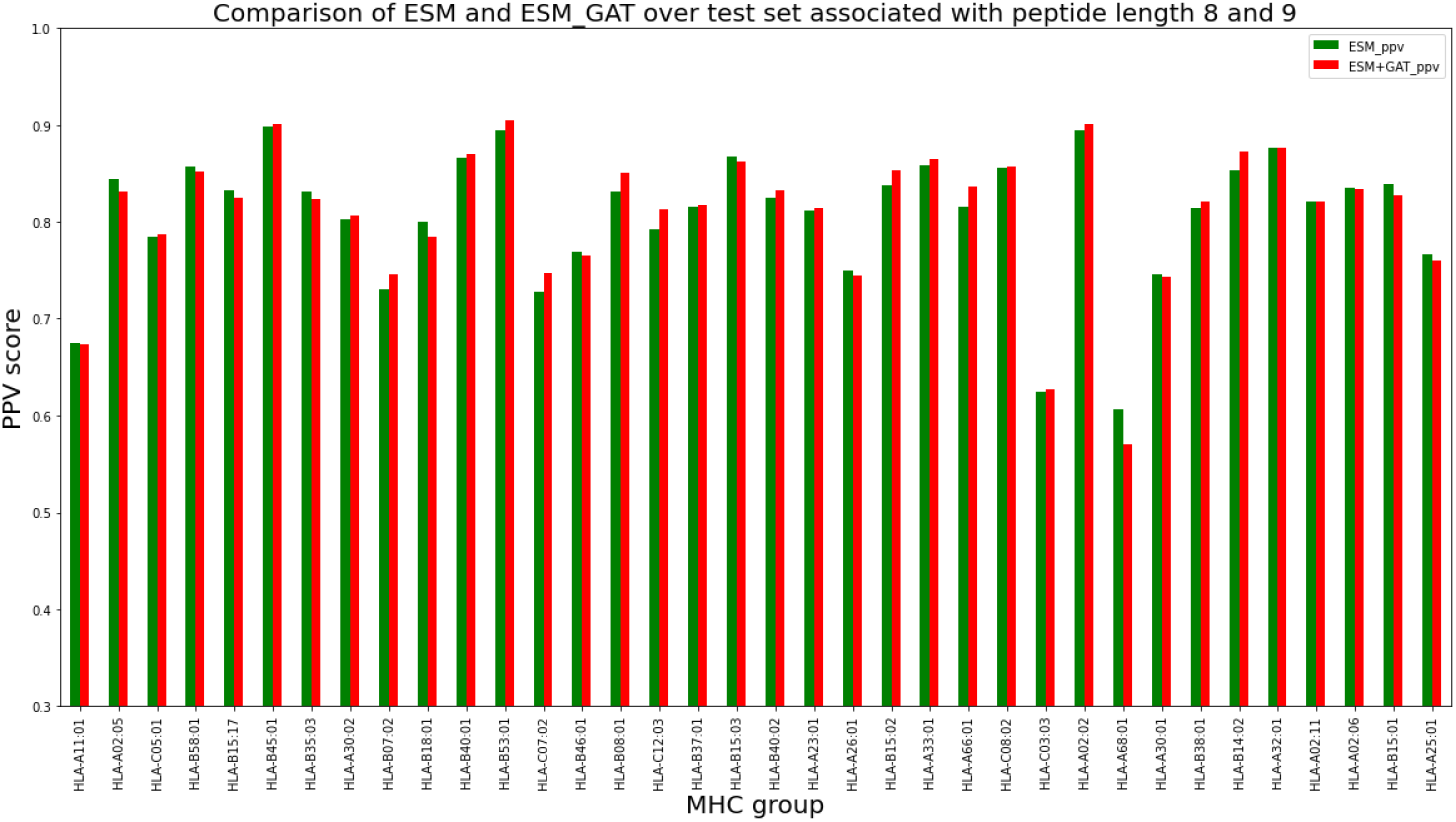
PPV Comparison (hit-decoy ratio: 1:19) of ESM fine-tuning method versus ESM-GAT over the training set with peptide length 8 and 9 and test set with peptide length 8 and 9.

## 4 Conclusion

Predicting peptides that bind to the major histocompatibility complex (MHC) Class I is an important problem in studying the immune system response and a plethora of approaches have been developed to tackle this problem. Among these, the most recent version of NetMHCpan server (NetMHCpan 4.1) [2] has been shown to achieve state-of-the-art performance. NetMHCpan 4.1 is developed based on training a shallow neural network, which, according to [2], outperforms other methods such as MHCflurry [47], MHCFlurry-EL, and MixMHCpred [44]. A number of recent works have focused on using protein language models in MHC-peptide binding problems. Protein language models developed based on deep learning approaches, such as attention-based transformer models, have shown significant progress towards solving a number of challenging problems in biology, most importantly, protein structure prediction [48]. BERTMHC [35] and ImmunoBERT [36] for the first time applied the pre-trained protein language models in MHC-peptide binding problems. Both methods used a relatively small pre-trained model (TAPE [26] which was trained with 31 million protein sequences); currently, there are substantially larger and more informative models such as ESM1b

[34] and ProtTrans [32] which are trained on more than 250 million protein sequences. BERTMHC was trained for MHC Class II and ImmunoBERT for MHC Class I; The focus of ImmunoBERT was on the interpretation of their model’s architecture and a fair comparison of the performance of their model with other works was not possible due to having different training sets. In the work reported in this paper we focus on MHC Class I peptide binding prediction by developing an approach based on a large pre-trained protein language model, ESM1b [34]; we follow two fine-tuning approaches using a soft-max layer and Graph Attention Transformer (GAT). In order to have a fair comparison, we train our model using the same training set used by NetMHCpan 4.1 [2] and evaluate our model using the same test set. We show, using the standard metrics in this area, that our model outperforms NetMHCpan 4.1 in 35 test sets out of 36. Since having the same training set is critical to compare different models, we did not compare our model directly with other works such as MHCflurry [4] given our different training sets. As reported, adding Graph Attention Network (GAT) to the ESM1b network, improved the ability of the model to predict peptides with lengths different from those considered in the training set; this feature is expected to be beneficial for training models beyond MHC Type I. Implementing a server based on our trained model is in progress which will be added to the Cluspro servers([49] [50] [51]).

## Acknowledgements

This work was supported in part by the National Institutes of Health grants R01 GM135930, RM1135136, R01GM140098, by the Boston University Clinical and Translational Science Award (CTSA) under NIH/NCATS grant UL54 TR004130; by the National Science Foundation grants IIS-1914792, DMS-1664644, DMS-2054251 and CNS-1645681; and by the Office of Naval Research grant N00014-19-1-2571.

## Footnotes

* A token is a string of contiguous characters between two spaces, or between a space and punctuation marks.

